# Effect of tributyltin chloride (TBT-Cl) exposure on expression of *HSP90β1* in the river pufferfish (*Takifugu obscurus*): evidences for its immunologic function involving in exploring process

**DOI:** 10.1101/279331

**Authors:** Xu Dong-po, Fang Di-an, Zhao Chang-sheng, Jiang Shu-lun, Hu Hao-yuan

## Abstract

*HSP90β1* (known as GP96) is a vital endoplasmic reticulum depended chaperonin among the HSPs family. It plays important roles in regulating the growth, development, differentiation, and apoptosis of cells. Furthermore, it always processes and presents antigen of the tumor and keeps balance for the intracellular environment. In the present study, we explored the effect of tributyltin chloride (TBT-Cl) exposure on *HSP90β1* expression in river pufferfish, *Takifugu obscurus*. The full length of *To-HSP90β1* was gained with 2775 bp in length, an ORF obtained with 2412 bp encoding an 803 aa polypeptide. The phylogenetic tree was constructed and showed the close relationship to other fish species. The *HSP90β1* mRNA transcript was expressed in all tissues investigated. After the acute and chronic exposure of TBT-Cl, the mRNA level of *To-HSP90β1* significantly up-regulated in tissues of liver and gill. Moreover, the histochemistry study indicated the injury degree of TBT-Cl on liver and gill. Immunohistochemistry (IHC) staining results implied the cytoplasm reorganization after TBT-Cl stress and the function of immunoregulation for *To-HSP90β1*. All the results indicated that *HSP90β1* may involve in the resistance to the invasion of TBT-Cl for keeping autoimmune homeostasis.

## 1. Introduction

Heat shock proteins (HSPs) are a series of special proteins which could generate and be activated in the environment of heat stress and other biological stress (Ritossa, 1962). HSPs are also called stress proteins (SPs) due to taking part in regulating other stress responses such as oxidative stress, heat, infection, toxicosis and so on (Erlejman et al., 2014a; Sørensen et al., 2003). It has been proved that HSPs have imperative roles in inhibiting protein aggregation, helping in folding the nascent proteins, and are considered to protect cells against oxidative stress, which are protection system to defend the organisms from harmful stress by preventing their reversible loss of vital proteins and facilitating their subsequent regeneration (Fu et al., 2011; Jiang et al., 2012; Parsell and Lindquist, 1993). HSPs derived from cancer cells or cells of viral infection could cause protective immunity, and their peptide-binding characters for specific vaccination served a potential approach to resisting the aggression of cancer and infectious diseases (Udono and Srivastava, 1993; Udono and Srivastava, 1994). HSPs are a cluster of highly conserved molecular chaperones which were ubiquitously expressed in tissues. They are segmented into distinct multigenic families, like HSP110, HSP90, HSP70, HSP60, HSP40 and other small HSPs. Among them, HSP90 is often found in a constitutive dimmer, which participates in controlling multiple regulatory pathways such as stress defense, hormone signaling, cell cycle control and apoptosis (Rajeshkumar et al., 2013). In *Crassostrea hongkongensis*, HSP90 plays a vital role in response to both osmotic stress and bacterial invasion (Fu et al., 2011). For many fish species, the HSP90 have been connected to cytoprotection and cell survival (Csermely et al., 1998; Smith et al., 2015), performing a protective and inducible role (Xu et al., 2014; Zhang et al., 2015). HSP90 in the liver was also found induced by ammonia stress, indicating that this kind of protein hammered at protecting body from oxidative stress and apoptosis (Cheng et al., 2015). It’s worth noting that *HSP90β1* (GP96), a subtype of HSP90 members, associated with major histocompatibility complex (MHC) class I molecule, which indicated it might be involved in immune response (Suto and Srivastava, 1995). The mRNA level of *HSP90β1* preferentially expressed in hepatocellular carcinoma and significantly increased in hepatoma cell line. Its expression had a down-regulation when the oncocytes differentiation was inducted by sodium butyrate. This indicated that *HSP90β1* had correlations with occurrence and development of cancer and cell differentiation (Cai et al., 1993; Heike et al., 2000a). Studies had shown that some stress factors made increases of expression level for GP96, in the meanwhile, its immunogenicity of GP96 was also aggrandized and rose with the expression level increasing (Dai et al., 2003). In the process of autoimmunity, the cell surface expression level of an endoplasmic reticulum (ER)-dependent GP96 initiated systemic autoimmune diseases in the body (Liu et al., 2003). Totally, *HSP90β1* exert great effects to raise the body immunity. Its special role in the study of anti-neoplastic immunity had been a hot topic for clinical immunotherapy (Conrad and Nestle, 2003; Mansour and Ronald, 2004).

Tributyltin chloride (TBT-Cl), is one of the most representative chemical compounds of Tributyltin (TBT). In view of its fatal toxicity to hydrobios, TBT was severed as a threat to water security (Antizar-Ladislao, 2008; Organization, 2001). TBT residual in water from various channels had become a noticeable problem, which made TBT contamination of aquatic ecosystems (Antizar-Ladislao, 2008; Tessier et al., 2007). TBT induced imposex in mollusks and fishes, which suggested that this toxic substance exerting a force on aquatic animal gonad function (Matthiessen, 2008; Mcallister and Kime, 2003; Nakayama et al., 2004; Shimasaki et al., 2003). Furthermore, TBT was found to be an inducer in the course of accumulation of adipose and altered fatty acid levels in male and female *Marisa cornuarietis* (Inadera and Shimomura, 2005; Janer et al., 2007; Meador et al., 2011). In zebrafish, TBT indeed altered multiple and complex activities of mRNA level in lipid metabolism and cell damage, which implied that underlying molecular mechanism of TBT on hepatic steatosis (Zhang et al., 2016).

*Takifugu obscures*, commonly known as river pufferfish, is an anadromous fish and an economic species, and studies on pufferfish aquaculture and its ecological environment have been a hot topic in the meanwhile (Kai et al., 2005; Van, 2004; Yamanoue et al., 2009). The pufferfish are important and scarce sources at the lower reaches of the Yangtze River and the river mouth area in China. As is mainly used as a biocide in antifouling agents applied to ships to prevent attachment of mollusks and hydrophyte (Antizar-Ladislao, 2008), the pulotions of TBT-Cl residuary in Yangtze River could exert an influence to its development of pufferfish. *T. obscures* was always chosen as a model to explore its adaptive and resisting mechanisms when exposed to different kinds of environmental stress factors (Kato et al., 2005; Kim et al., 2010a). However, the physiological function of the pufferfish under the exposure of TBT-Cl keeps unclear. It’s attractive to us that studying for its mechanism of *T. obscures* exposed to TBT-Cl may have a profound meaning.

In this paper, *HSP90β1* gene in *T. obscures* from databases of transcriptome sequencing was characterized by bioinformatic analysis, and the phylogenetic tree was constructed based on HSP90 sequences of other species. Tissues expressions of this gene were detected by quantitative real-time PCR (qPCR) method. After exposing to different concentrations of TBT-Cl in the acute and chronic experiment, the *HSP90β1* mRNA level was checked through qPCR. The histochemistry and IHC test were performed to verify the damaging effect of TBT-Cl to the pufferfish. This study may supply a deeper understanding of the unique function of *HSP90β1* in the course of fighting with the adverse effect of TBT-Cl and explain the conceivable mechanism in immunoreaction.

## 2. Materials and methods

### 2.1. Animals

*T. obscurus* with an average length of 10 ± 1.5 cm and an average weight of 25.1 ± 2.23 g were obtained from the aquaculture base in Freshwater Fisheries Research Center (FFRC, Wuxi, China). The pufferfish were kept in 100-L cylindrical opaque polypropylene aquaria and supplied with commercial feed twice a day at regular intervals. After at least 7-day acclimation, robust animals were chosen until 24 h-feeding before the experimental treatments. The water was exposed to air for a week to remove chlorine. During the experiment, the temperature kept at 26 ± 2 °C, the dissolved oxygen and pH maintained at 7.93 ± 0.45 mg/L and 7.83 ± 0.12, respectively. All the operations to the pufferfish were carried out in strict accordance with the recommendation in the criterion for the care and use of laboratory animals.

### 2.2. TBT-Cl exposure and sampling

Healthy fish were randomly chosen and divided into four groups. On the basis of 96 h acute toxicity experiment (96 h-LC_50_ = 19.62 μg/L), the pufferfish were exposed to three kind of concentrations of TBT-Cl (10% 96 h-LC_50_, 20% 96 h-LC_50_ and 50% 96 h-LC_50_) and the DMSO solution (V (DMSO): V (water) = 1%). Ten individuals were put into a group randomly. After the exposure, at the time point of 96 h, fish were collected (n = 6) and anesthetized in diluted tricaine methanesulfonate (MS-222, Sigma, USA) at the concentration of 100 mg/L. The fish were put on the ice and sampled with blood, liver, gill, heart, muscle, stomach, intestine, kidney, spleen and brain.

For the chronic toxicity experiment, the treatment group (900 ng/L of TBT-Cl) and a control group (DMSO group) were set. Each group had six repetitions (n = 6), 10 fish were in each repetition. The experimental period was 30 days, and sampling was performed every 10 days. The fish was collected in each repetition randomly. Blood was extracted and the brain, liver, gill, stomach, intestine, heart, muscle, kidney and spleen were sampled.

After 30-d exposure experiment, the recovery test was followed. The water in all groups was changed to aerated tap-water. The period was 30 days and fish were collected every 15 days. Six animals were gained in each group randomly (n = 6). After normal saline wash, the fish put on ice were rapidly sampled and its blood, brain, liver, gill, heart, muscle, stomach, intestine, kidney and spleen were sampled. All the serum and tissues were snap-frozen in liquid nitrogen after labeled and stored at −80 °C for later assay.

### 2.3. Total RNA extraction and cDNA preparing

Total RNA was isolated from the harvested pufferfish tissue using Trizol reagent (Invitrogen, USA) according to the manufacturer’s instruction, then dissolved in DEPC (diethylpyrocarbonate)-treated water and stored at −80° C. The cDNA template was prepared containing 2 μg total RNA by reverse transcription reaction by a PrimeScript^TM^ RT reagent K it with gDNA Eraser (Perfect Real Time) (TaKaRa, Japan) following the manufacturer’s protocol. The concentration and quality of RNA and DNA products were measured by spectrophotometry (absorbance at 260 nm) and agarose gel electrophoresis, respectively.

### 2.4. The full-length cloning and phylogenetic analysis

Target sequences of cDNAs encoding *HSP90β1* were obtained from the libraries of transcriptome sequencing (unpublished data). The specific primer for *HSP90β1* was designed using Primer Premier 5 (*To-HSP90β1*-S: TGGTGGGAGCGGTGGCTTGTCAGTCCTCTTGT; *To-HSP90β1*-A: AGAACCACAGTGGAGCTGGAACTCTCAGAC). The full-length template for cloning *To*-*HSP90β1* was verified by PCR amplification. Its product was determined by agarose gel electrophoresis. The biological sequence obtained was analyzed by BLAST <u>(https://blast.ncbi.nlm.nih.gov/Blast.cgi)</u> in NCBI (Mcginnis and Madden, 2004). The program ClustalW2 was used to perform multiple sequence alignment (Chenna et al., 2003; Thompson et al., 2002). The phylogenic tree was constructed with MEGA 6.0 through a neighbor-joining (NJ) algorithm based on the deduced amino acid sequence of HSP90 for some other species (Kelly et al., 2006; Yu et al., 2015).

### 2.5. qPCR detection of tissues expression patterns

The cDNA sample (n = 6) collected from blood, liver, stomach, intestine, gill, heart, muscle, kidney, spleen and brain above were all performed to determine their expressions by qPCR. The primer of *To*-*HSP90β*1 was designed (RT-*HSP90β1*-F: CCCTGGAGAAGGACTTTGAGC, RT-*HSP90β1*-R: GGGGTGTTTGGGGTTGATTT). *β-actin* (RT-*β-actin*-F: AGAGGGAAATCGTGCGTGAC, RT-*β-actin*-R: CAAGGAAGGATGGCTGGAAG) in *T. obscurus* (GeneBank accession number: EU871643) was measured as the internal control to normalize the level of qPCR results. Before beginning the qPCR program, the specificity and efficiency of primers were tested. The reaction system was carried out in a total volume of 20 μL, including 10 μL of SYBR *Premix Ex Taq* II (TaKaRa, Japan), 2 μL of cDNA template (80 ng total RNA), 1.6 μL of both sense and anti-sense primers (10 μM), 0.4 μL of ROX Reference Dye (50 ×) and 6 μL of PCR-grade water, which carried out in triplicate. Two-step PCR program was performed, which containing of 1 cycle of 94 °C for 35 s, 40 cycles of 95 °C for 10 s, 59 °C for 30 s, followed by 1 cycle of 95 °C for 15 s, 60 °C for 60 s and 95 °C for 30 s. qPCR results were calculated by using ABI StepOnePlus Real-Time PCR software (Applied Biosystems, USA) with 2^−ΔΔCt^ methods (Schmittgen and Livak, 2008).

### 2.6. The qPCR detection of *To-HSP90β1*

qPCR was performed to determine the expression level of *HSP90β1* in the liver treated with TBT-Cl at timepoint 96 h under the concentration of 0, 10%, 20% and 50% of 96 h-LC_50_ TBT-Cl using the gene-specific primers (see in Section 2.5). It was also detected in chronic toxicity and recovery test. The reaction process referred to the operation Section above. The qPCR of each sample carried out in triplicate (n = 3). Besides, the expression of *To-β-actin* was measured and used as the internal control to normalize the results of qPCR analysis.

### 2.7. Preparation of paraffin section

The fresh gill and liver tissues were fastened with 10% neutral formalin for 24 h. The dehydration was reached with the different concentrations of ethanol. And, the waxed tissues were buried in the embedding machine and the dressed block was sliced, flattened then baked at 65 °C.

The paraffin section was dewaxed, washed, and stained with cold hematoxylin for 8 min. After washed by freshwater, the section was stained with eosin stain for 3 min. When dehydrated by ethanol and dimethylbenzene, the histological section was sealed by neutral gum.

### 2.8. The anti-HSP90β1 antibody preparation

The synthetic polypeptide and monoclonal antibody were enforced commercially by Abcam (Abcam, England). In short, HSP90β with a synthetic C-terminal peptide (EDASRMEEVD) combined with keyhole limpet hemocyanin was emulsified with complete Freund adjuvant for the first immunization and incomplete Freund adjuvant for the second to fourth immunizations and was injected into a New Zealand rabbit at one-month interval. Before the fourth immunization, its serums of the rabbit were sampled. An increase in antibody titers against the peptide was verified by enzyme-linked immunosorbent assay (ELISA).

### 2.9. Immunohistochemistry

Paraffin sections were used for IHC analyses. Livers and gills at different sampling stages were taken out and fixed in 0.01 M phosphate-buffered saline (PBS) containing 4% of paraformaldehyde at 4°C for above 6 h. After PBS washing for three times, the samples were dehydrated in 30% saccharose-PBS solutions for 4 h at room temperature and then embedded in organ optimal cutting temperature compound (Sakure, USA). Standard sections of 8 μm in thickness were taken using a microtome (Leica, Germany). IHC was put into effect according to the modified manual (Multhoff, 2007). Briefly, the sections were washed for three times with 0.01 M PBS for 15 min each wash. Sections were soaked with 0.01 M citric acid buffer (pH 6.0) which contained 0.1% of Tween 20 and autoclaved for 5 min. The sections were blocked in PBS (pH 7.4) and incubated with anti-HSP90β (1:200) overnight at 4°C. Then the sections were washed three times with 0.01 M PBS for 10 min each wash. Subsequently, the tissue sections were incubated with secondary antibody goat anti-rabbit IgG conjugated with horseradish peroxidase for 30 min, and then rinsed three times for 5 min each wash with PBS. Immunoreactive signals were observed by diaminobenzidine (Sigma, Japan) as the substrate. The sections were counterstained with hematoxylin-eosin (HE). Incubated buffers preimmune rabbit serum and the blocking solution were also used to treat organ sections as the negative control.

### 2.10. Statistical analysis

IBM SPSS Statistics 19 (Chicago, IL, USA) was used and the significant difference was shown by student’s t-test and one-way ANOVA (one-way analysis of variance) by the mean of comparing means between samples in the process of data analysis. The P value set below 0.05 was considered to be statistically significant (signed as a, b, c or d). The results we got were presented as means ± SD (standard deviation).

## 3. Results

### 3.1. The characterization and phylogenetic analysis

After exploring the transcriptome libraries, a member of *To*-*HSP90* was determined: *To-HSP90β1* (GeneBank accession number: MG597234). The full length was obtained as 2775 bp in length, with the ORF (open reading frame) of 2412 bp. Besides, it contained a 112 bp of 5’-UTR (untranslated region) and a 251 bp of 3’-UTR (Fig. 1). *ToHSP90β1* ORF sequence encoded an 803 aa polypeptide, the molecular mass of 92.36 kDa and the pI of 4.72. After the InterPro sequence search, in *HSP90β1*, four homologous superfamilies were found: two Histidine kinase/HSP90-like ATPase superfamilies (position 77-299 aa and 338-375 aa), a Ribosomal protein S5 domain 2-type fold (position 345-600 aa) and an HSP90, C-terminal domain (position 624-749 aa). Several feasible functional domains were detected in To-HSP90β1 using Motif Scan tool (https://myhits.isb-sib.ch/cgi-bin/motif_scan) (Liu et al., 2002; Periannan et al., 2012). The To-HSP90β1 sequence consisted of an Amidation site, four Asn_glycosylation sites, three cAMP_phospho_site, seventeen CK2_phospho_site, seven MYRISTYL sites, sixteen PKC_phospho_site, a RGD site, three TYR_phospho_site, an ER_Target domain, a Heat shock hsp90 proteins family signature domain, an aspartic acid-rich region profile, a ELM2 domain profile, a Glutamic acid-rich region profile, a Histidine kinase domain profile, a Bipartite nuclear localization signal profile, a Protein prenyltransferases alpha subunit repeat profile, a Histidine kinase-, DNA gyrase B-, and HSP90-like ATPase and an Octapeptide repeat. The sequence also contained an HSP90 domain and a chaperone protein htpG signature (marked in Fig. 1), which showed that this protein really played specific roles due to its motifs.

**Fig. 1.**
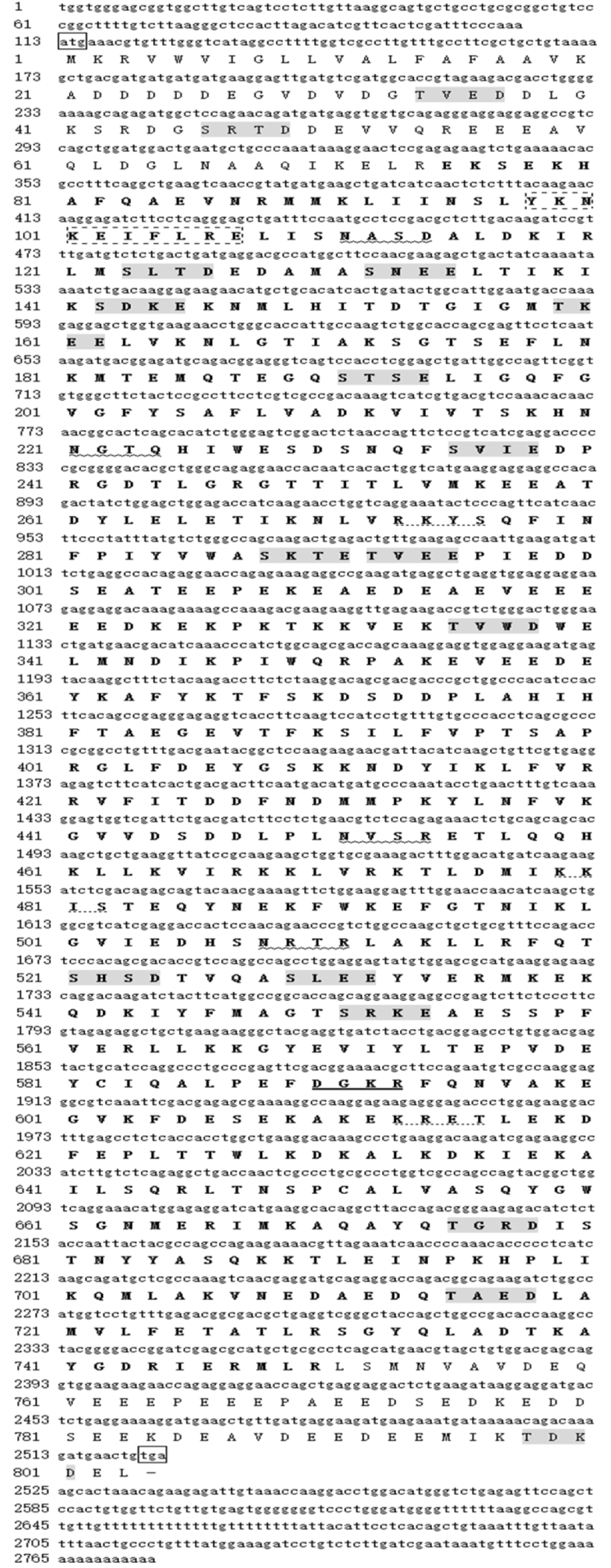
The start codon (ATG) and stop codon (TGA) are boxed with solid lines. The Amidation site is underlined with solid lines in bold, Asn_glycosylation sites are showed with wavy lines, the cAMP_phospho_site is underlined with dotted lines, the CK2_phospho_site is in light grey, the HSP90 domain is boxed with dotted lines and a chaperone protein htpG signature is in bold.

BLAST analysis revealed that *To-HSP90β1* shared high similarity with other HSP90s, including those from *Takifugu rubripes* HSP90β1 (99%), *Notothenia coriiceps* HSP90β1 (86%), *Lates calcarifer* HSP90β1 (86%), *Larimichthys crocea* HSP90β1 (87%), *Monopterus albus* HSP90β1 (86%), *Oreochromis niloticus* HSP90β1 (85%), *Paralichthys olivaceus* HSP90β1 (85%), *Oryzias latipes* HSP90β1 (84%), *Salmo salar* HSP90β1 (84%), *Oncorhynchus mykiss* HSP90β1 (83%), *Danio rerio* HSP90β1 (82%), *Rattus norveqicus* HSP90β1 (76%) and *Scylla paramamosain* HSP90 (72%). The *HSP90β1* and members of *HSP90* for other species were used to construct the phylogenetic tree by Clustal 1.81 and MEGA 6.0. The sequence *To-HSP90β1* was most closed the species of *T. rubripes HSP90β1*, which indicated a significant correlation of genetic relationship for these two fish in evolution. The NJ phylogenetic tree contained four distinct branches, where *T. obscurus* clustered with fish species especially for anadromous fish, with mammals and birds formed a different cluster. In addition, the conservational and phylogenic clustering of eukaryote HSP90 sequence is consistent with eukaryotic classification (Fig. 2).

**Fig. 2.**
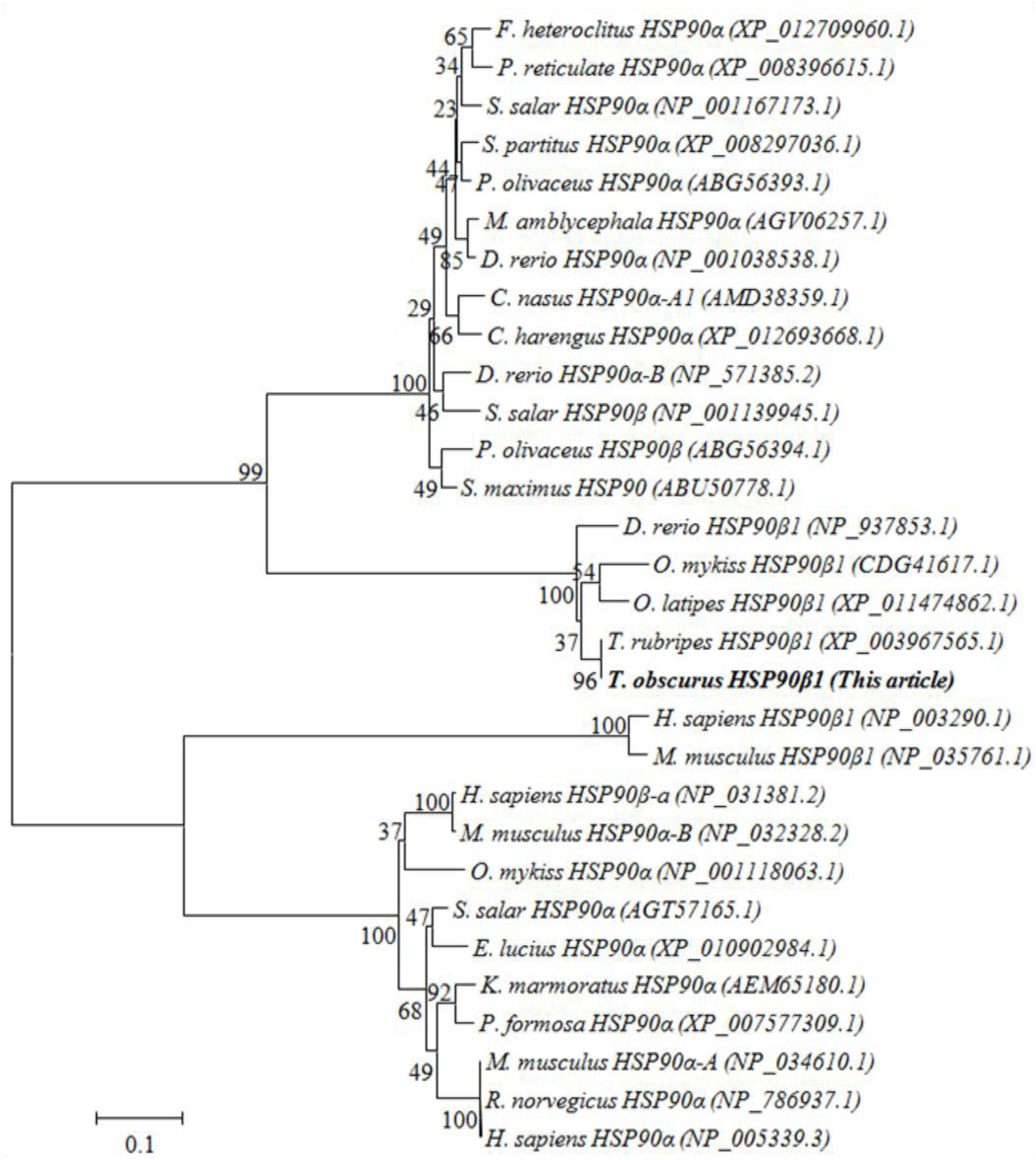
The phylogenetic tree analysis of HSP90. The phylogenetic tree was constructed on the base of a series of *HSP90β1* proteins from different species. *T. obscurus HSP90β1* is shown in bold. The analysis is based on proteins from the above mentioned species. Phylogenetic tree constructed by the MEGA 6.0 program by the neighbor-joining (NJ) distance method. The statistical robustness of the tree was estimated by bootstrapping with 1000 replicates. Bootstrap values were indicated by genetic distance. The putative protein GenBank accession number is shown in parentheses.

### 3.2. *To-HSP90β1* expression pattern in tissues and after TBT-Cl exposure

The pattern of *HSP90β1* in river pufferfish was ubiquitously expressed in all the detected tissues: blood, heart, gill, liver, stomach, intestine, muscle, brain, kidney and spleen. In Fig. 3, the liver and gill tissues had the most abundant amount of *HSP90β1* transcript, which was obviously higher than other tissues.

**Fig. 3.**
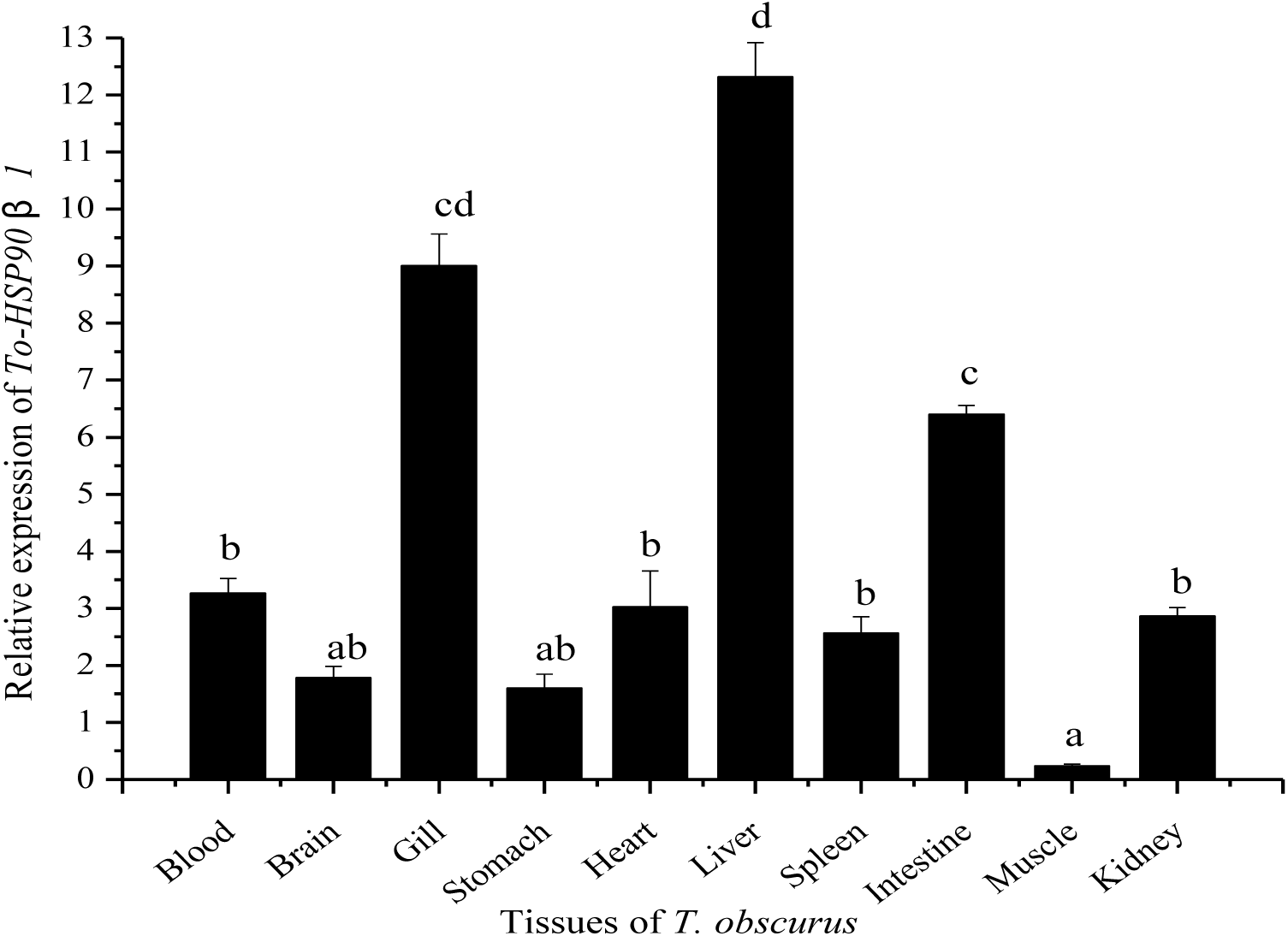
Tissues expressions of *To*-*HSP90β1*. The relative *To-HSP90β1* mRNA levels derived from ten tissues for six individuals in each group were calculated by the 2^−ΔΔCt^ method. Expression levels are normalized using *To-β-actin*. Vertical bar shows the mean ± SD (n = 6). Significant differences (P < 0.05) are expressed with superscript letters (a, b, c and d, respectively, a < b < c < d).

*To-HSP90β1* expression after TBT-Cl was validated by qPCR method. In Fig. 4A, its *To-HSP90β1* expression patterns were all significantly up-regulated with the increase of the concentration of TBT-Cl both in liver and gill. In gill, its expression was sharply up-regulated at 10% LC_50_-96 h of TBT-Cl and then increased until at 50% LC_50_-96 h of TBT-Cl, while had a fluctuation at 20% of the concentration. However, the mRNA level in liver was relatively gentle. Broadly speaking, the expressional level in gill was higher than that in the liver. In the chronic experiment, the mRNA level rose prominently from 0 d to 20 d and down-regulated extremely at 30 d for gill sample. *To-HSP90β1* expression was smooth at recover stage in the gill. Nevertheless, its expression in liver kept a low level. In the whole processes, the 20 d-sampled group in chronic exposure stage at a lower concentration of TBT-Cl had the highest level of *To-HSP90β1* than other groups in the acute and chronic test. The entire expression pattern above showed that *To-HSP90β1* could react significantly to the effect of TBT-Cl.

**Fig. 4.**
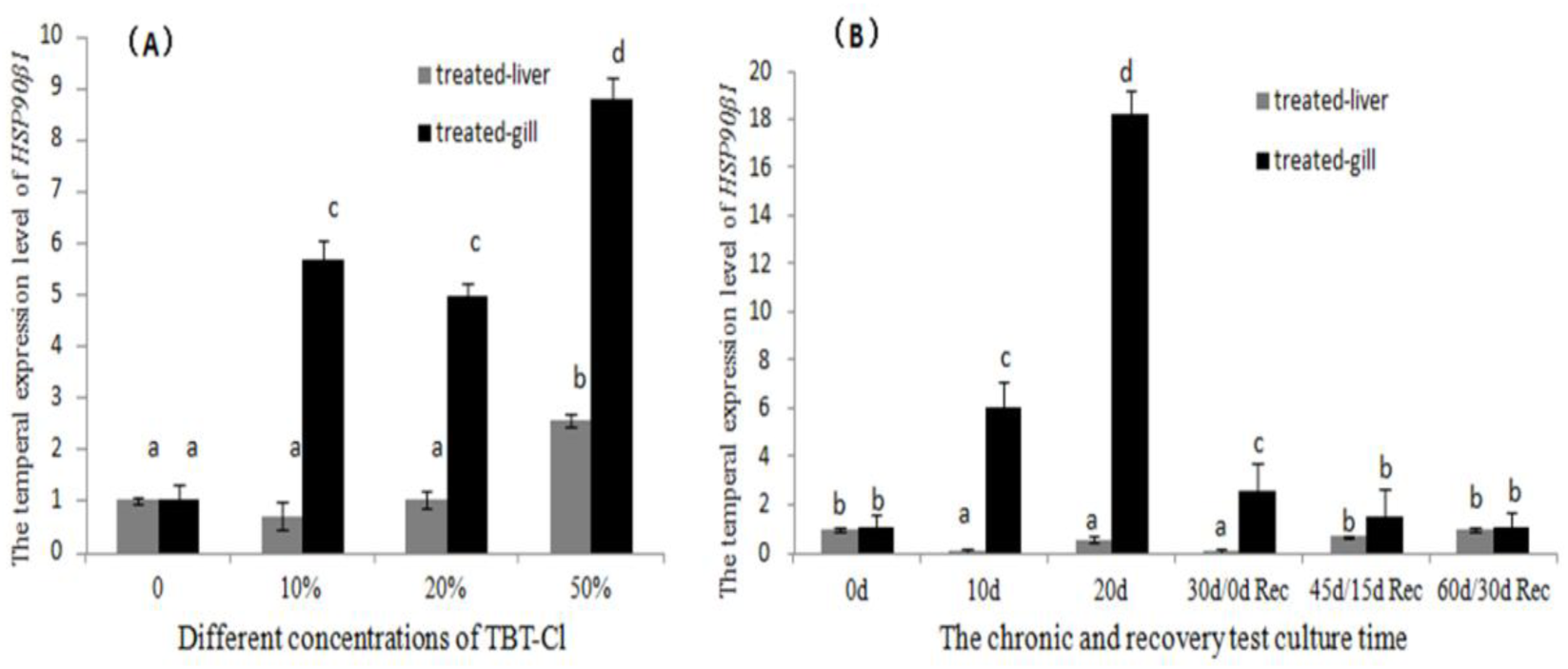
The mRNA levels of *To-HSP90β1*. (A) The pufferfish were exposed to four concentrations of TBT-Cl (0, 10%, 20% and 50% LC_50_-96 h). (B) The pufferfish were exposed to 900 ng/L of TBT-Cl. 30 d was designated as 0 d for recover. The samples were collected in sextuplicate (n = 6). Significant differences (P < 0.05) are presented with different superscript letters (a, b, c and d, respectively, a < b < c < d).

### 3.3. Histochemistry and immunohistochemistry

In order to verify its function of *HSP90β1* after TBT-Cl exposure, the histochemistry and immunohistochemistry were performed in liver and gill. Fig. 5 (L1) showed a normal liver with even clear cells and pancreas. In L2, the tissues began to show vacuolation at the 10% LC50-96 h of TBT-Cl. With the concentration of LC50-96 h of TBT-Cl added to 20%, the tissues appeared to cellular edema and indistinct cytoplasmic borders in L3. In Fig. 5 (L4), the liver cells were necrotic under a high dosage of 50% of LC50-96 h of TBT-Cl. Moreover, the ducts in the pancreas are thickened. As shown in L5, the liver had no signals while the signals of *HSP90β1* in the pictures from L6 to L8 with the rise of TBT-Cl concentration became more and more obviously. The target protein of pancreatic epithelial cells increased, and the target protein decreased in hepatocytes and hepatic sinuses with the rise of degree of exposure. However, when it came to gill, these tissues in control group had structural integrity in Fig. 5 (G1). In G2, intercellular space of the gill became small, and the gill raker was anomalous; the gill filaments were wizened and deformative in G3; some epithelial cells exfoliated and myxocytes were swollen in the gill branch leaves under a 50% of LC_50_-96 h TBT-Cl in G4. For IHC in gills, the control group G5 showed no signals. After the exposure to 10% of LC_50_-96 h of TBT-Cl, the signal of *HSP90β1* enhanced evidently in gill filaments in G6. From G7 to G8, the mRNA level increased and its signal strengthened further. Myxocytes became intumescent in 20% of LC_50_-96 h of TBT-Cl while the myxocytes vanished in 50% LC_50_-96 h of TBT-Cl exposed liver.

**Fig. 5.**
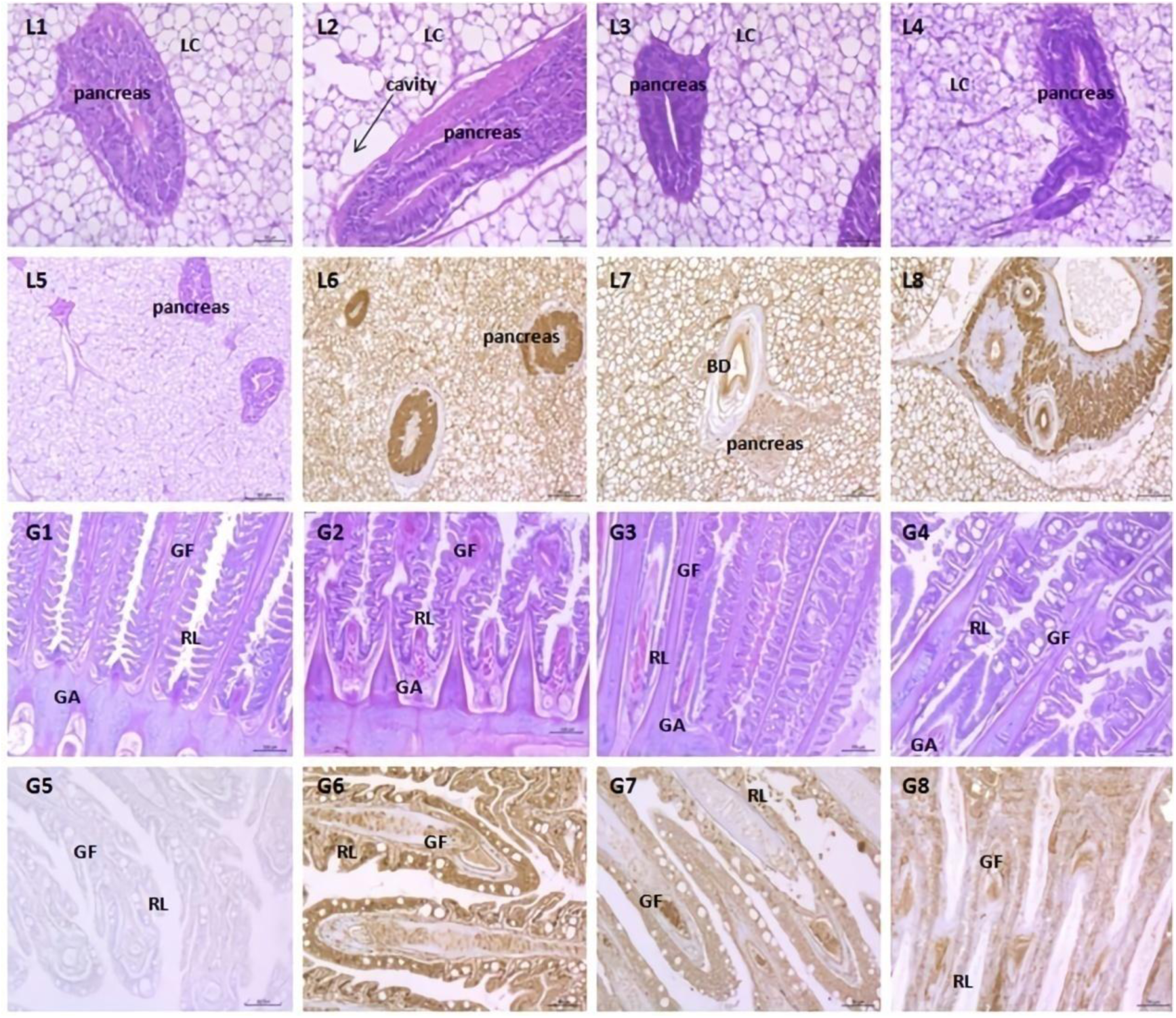
Histochemistry (L1, L2, L3, L4, G1, G2, G3 and G4) and immunohistochemistry (L5, L6, L7, L8, G5, G6, G7 and G8) for *HSP90β1* in liver and gill. L1: The control group of HE stain in liver; L2, L3 and L4: HE stained with 10%, 20%, and 50% of LC_50_-96 h TBT-Cl exposed in liver, respectively; L5: IHC for control liver, L6, L7, and L8: IHC for 10%, 20%, and 50% of LC_50_-96 h in TBT-Cl exposed liver, respectively. G1: The control group of HE stain in liver; G2, G3 and G4: HE stained with 10%, 20%, and 50% of LC_50_-96 h TBT-Cl exposed in liver, respectively; G5: IHC for control liver, G6, G7, and G8: IHC for 10%, 20%, and 50% of LC_50_-96 h in TBT-Cl exposed liver, respectively. LC: liver cell; BD: bile duct; GF: gill filament; GA: gill arch; RL: branch leaf.

## 4. Discussion

As a stress-sensitive molecular chaperone, HSPs played essential roles in a series of metabolism processes (Xie et al., 2015). A lot of treatments on cells activated the expression of HSP genes (Song et al., 2016). *HSP90β1*, a special actor in stress response, is an ER-enriched distributed protein which participates in associating with neonatal or abnormal proteins, assisting repair and thermo-resistance of cells (Berwin et al., 2002). Moreover, its overexpression on the cell surface was deemed to play roles in immunology relative reactions (Liu et al., 2003). In our present study, the full-length cDNA sequence encoding *HSP90β1* indicated that it has two signature sequences motifs consistent with other HSP90 family proteins: a stress-induced protein motif and a terminal C-terminal domain. A special ER-target domain is also included in the sequence. This indicates likely cytosolic localization of *HSP90β1* and suggests that *HSP90β1* contains the typical conserved structural features of other eukaryotic cytoplasmic HSP90s. Moreover, the C-terminal TDKDEL characteristic of cytosolic HSP members that mediates inter-domain communication and peptide-binding capacity (Stetler et al., 2010), as well as other additional important residues involved in ATP hydrolysis, ATP binding and ATPase activity, interdomain interaction and phosphorylation by casein kinase II were also detected suggesting that both *HSP90β1* genes are functional. Obviously, this sequence included some domains and relative specific motifs, which indicates that it is conserved in evolution and involved in the relative biological process. The phylogenetic tree constructed on the basis of HSP90 sequences showed that *To-HSP90β1* had the closest genetic relatives to the protein of *Takifugu rubripes*. It indicated that *To-HSP90β1* clustered in a most typical *HSP90β1* family of other species, which shows a conserved domain of functional structure in evolution. The neighbor-joining phylogenetic tree reveals a high degree of conservation in the HSP90 multigene family during evolution (Yeyati and van Heyningen, 2008). The phylogenic tree indicates an early origin for the HSP90 ortholog in eukaryote evolution (Erlejman et al., 2014b).

In order to clarify its distributions of *To-HSP90β1* in tissues, the pattern expressions were found strongest in liver and gill, indicating that the two immune-associated tissues may involve in the main resistance to the invasion of infaust environmental factors. It might guide us to the following exploration of *To-HSP90β1* function exposed to TBT-Cl.

To make certain the special response of this protein in the pufferfish after the invasion of TBT-Cl, the fish were exposed to different concentrations of TBT-Cl in the acute test. *To-HSP90β1* mRNA levels were significantly up-regulated at 96 h along with a rise of TBT-Cl concentrations in liver than that in gill tissue. It showed that TBT-Cl might activate the relative functional structure in *To-HSP90β1*. The increase of this protein response to the stimulation of toxic substance and it mobilized positive protective effect in homeostatic equilibrium. Interestingly, this gene in gill had a more distinct change, manifesting that the gill may be a crucial tissue for *To-HSP90β1*. In order to further study detailed functions of *To-HSP90β1* in pufferfish, chronic test and recovery experiment were enforced. Moreover, in the chronic toxicity treatment, *To-HSP90β1* showed a more serious rise during the first twenty days in gill, which indicated that this protein may take part in withstanding the impairment of TBT-Cl in the initial stage. Its sharp decline in the TBT-Cl-removed aquatic environment again displayed the effect of TBT-Cl to *To-HSP90β1*. After 30 d exposure, the pufferfish had adapted to toxicity stimulation and kept a moderate level of *HSP90β1* after a recover of damage to their bodies. *To-HSP90β1* may play essential roles in response to TBT-Cl exposure.

In order to verify its function to TBT-Cl exposure, the paraffin section of histochemistry and immunohistochemistry were performed. Normal liver and gill tissues of pufferfish showed moderate staining intensity for *To-HSP90β1*, while TBT-Cl exposure-tissues displaying significantly strong level signals and different degrees of tissue damage and cytopathy, which showed a tendency that the toxicity of soluble TBT-Cl in water induce *To-HSP90β1* to act as a resistant. It also indicated that *To-HSP90β1* may involve in the course of immunologic balance to reply to pessimal stimulation.

As an ER-located chaperone, *HSP90β1*, known as glucose-regulated protein 94 (GRP94 or GP96), took part in regulating cellular homeostasis and cancer biology (Heike et al., 2000b; Lammert et al., 1997; Melnick et al., 1994; Spee and Neefjes, 1997). *HSP90β1* was a central regulator in the folding of the protein and monitoring the activation of transmembrane ER stress sensors. The liver defense system may be triggered by the exposure of TBT-Cl, following by a liver pathological change (hydroncus even disruptive damage) in pufferfish. This signal was then transferred to the ER. Within the cell, especially in ER, *HSP90β1* acted as a luminal chaperone for protein recognition. On one hand, HSP40s, another chaperones in the process of folding and unfolding as well as translocation and degradation of proteins, share common substrates and interaction with HSP70s through binding to the latter N-terminal ATPase domain (Clare and Saibil, 2013; Goffin and Georgopoulos, 1998; Greene et al., 1998; Hernández et al., 2002; Johnson and Craig, 2001; Szabo et al., 1996). And ATP hydrolysis requires the involvement of nucleotide exchange factors (NEFs) mediating the subsequent binding of ATP which governs substrate release from HSP70s and sets back the HSP70 chaperone cycle (Brehmer et al., 2001; Harrison et al., 1997; Liberek et al., 1991). Moreover, HSP40s are implicated in the HSP90 chaperone pathway in conjunction with HSP70 thus cooperating in the folding of numerous substrate proteins in the cytosol of eukaryotes (Cintron and Toft, 2006). On the other hand, heavy-chain binding protein (BiP, GRP78, glucose-regulated protein 78) involved in proteins misfolded bind process and lead to proteins degradation through the proteasome in a process called ER-associated degradation (ERAD). Accumulation of misfolded proteins in the ER causes a stress and activates the unfolded protein response (UPR) signaling pathway. In certain severe situations, however, the protective mechanisms activated by the UPR are not sufficient to restore normal ER function and cells die by apoptosis (Määttänen et al., 2010; Naidoo, 2009; Stolz and Wolf, 2010). But, the activation of *HSP90β1* directly led to its increase of mRNA level. Then the degradation of its aim proteins and a series of cytoplasmic recombinations happened (Fig. 6). *HSP90β1* may play essential roles in this pathway. On the basis of the phenomenon we observed, an assumption was established: firstly, the exposure of TBT-Cl caused a reaction for non-specific and specific immunity, activating a series of inflammatory cytokines to resist the damage of this toxic chemical. Secondly, its toxicity of TBT-Cl induced the pressure for ER, and this energy might disequilibrate to the cellular environmental homeostasis. This disturbance in turn altered some functional molecular structures or chaperones mRNA levels, like *HSP90β1*, due to the existence of 5’-flanking region of *HSP90β1* contained ER-stress response elements (Nagahori et al., 2010). High level of TBT-Cl truly induced an increase of gp96 mRNA to react as a guard while chronic exposure triggered more dramatic rise of the mRNA level, which could conclude a relatively low concentration of TBT-Cl contributing to activation for *HSP90β1* in pufferfish. Interactions of ER-dependent components in a number of signal pathways regulated the body balance of *T. obscures*.

**Fig. 6.**
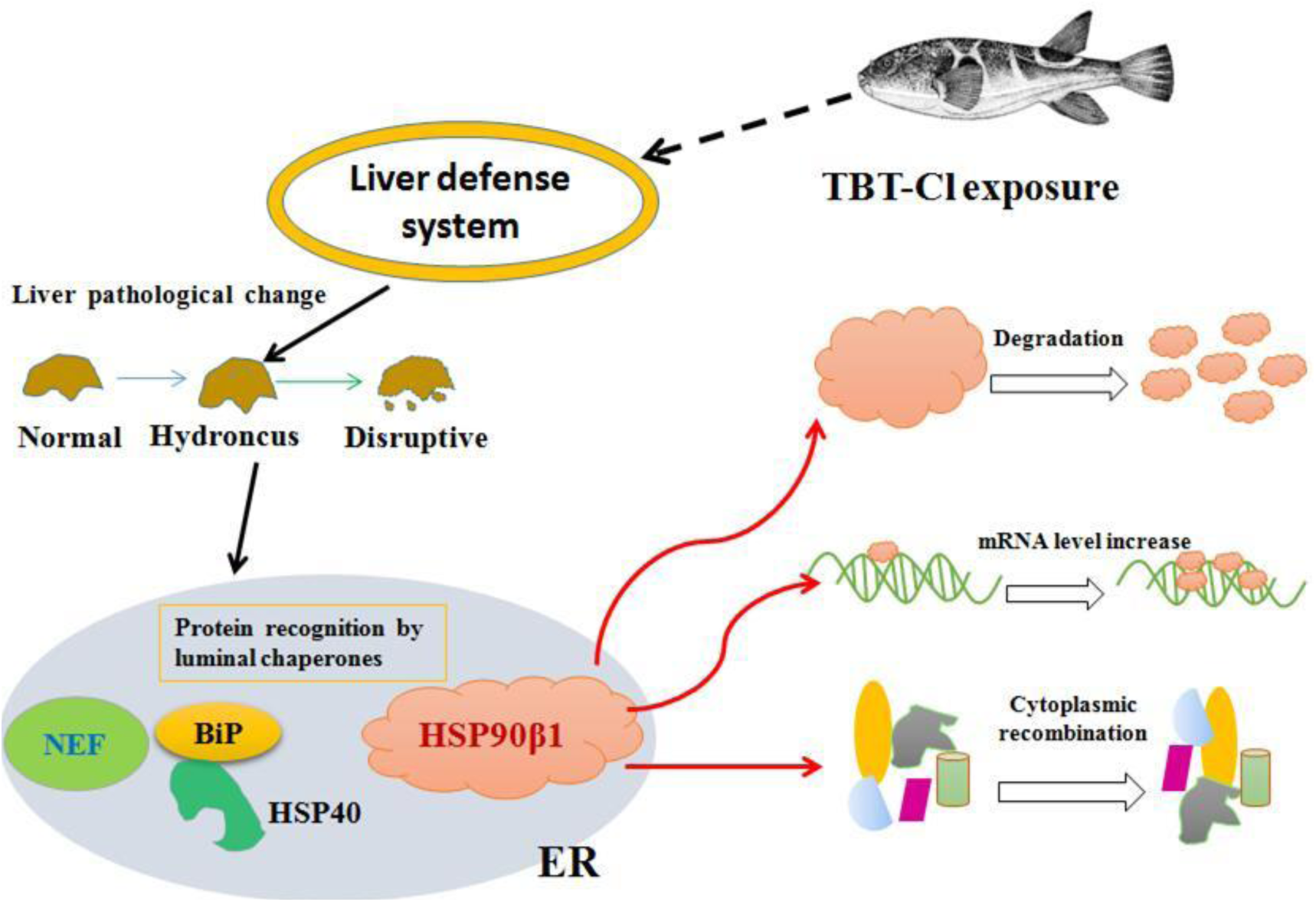
The possible immunologic injury regulation mechanism of *To-HSP90β1* involved in TBT-Cl exposure.

*T. obscures*, is often saw as a module to study the aquatic species coping with harmful factors (Ai et al., 2011; Kim et al., 2010b). When facing with TBT-Cl, its physical and functional changes accompanied. HSP90s, as important members to take part in unfolding, translocating and disintegrating proteins, took an important part in protecting cells against oxidative stress (And and Lindquist, 1993). *HSP90β1* was a chaperone of HSP90 family, showed mRNA level changes under the role of TBT-Cl in *T. obscures*. Following our study, the liver and gill were identified as vital tissues at the front lines to react to the invasion of TBT-Cl. The study we perform brought us a new outlook to regard glucose-regulated protein HSP90β1 facing with residual toxicant like TBT-Cl. It also developed our thought on high-yield fish culture and made the enlightenment to deeper study.

## Acknowledgments

This work was supported by funds from the National Infrastructure of Fishery Germplasm Resources (2017DKA3047-003) and the Fund from the Provincial Key Laboratory of Conservation and Utilization of Important Biological Resources in Anhui.

## Author Contributions

Xu Dong-po was responsible for data scoring and analyses, and writing the manuscript. Hu Hao-yuan conceived and designed the experiments. Fang Di-an, Zhao Chang-sheng and Jiang Shu-lun helped selecting the pufferfish tissues sample, RNA extraction and data analysis during manuscript preparation. All authors have read and approved the final manuscript.

